# Evaluation of potential stress marker genes during acclimation of the Patagonian *Lactiplantibacillus plantarum* UNQLp11 strain

**DOI:** 10.1101/2025.08.28.672453

**Authors:** Zerbino Antonella, Lannutti Lucas, Olguin Nair

## Abstract

In this study, we evaluated the expression of genes that may serve as markers of temperature- and ethanol-induced stress responses during the acclimation of *Lactiplantibacillus plantarum* strains of enological interest. The transcriptional responses of eight genes potentially related to stress adaptation were analyzed in two strains acclimated at 18□°C and 21□°C. Gene expression was assessed at the start (time zero) and after 48 hours of acclimation. Following acclimation, the cells were inoculated into sterile Pinot noir wine, and L-malic acid consumption was monitored over 19 days to determine whether acclimation temperature influenced cell survival and fermentation performance. Both strains exhibited similar L-malic acid consumption and survival patterns, although transcriptional responses varied depending on the temperature. Notably, *gapB, glmS*, and *rfbB* emerged as promising gene markers for future studies. These findings suggest that gene expression profiling of acclimated cells may support the selection of robust *Lpb. plantarum* strains for use as starter cultures in winemaking.

## Introduction

In wine, lactic acid bacteria (LAB) are responsible for malolactic fermentation (MLF), a decarboxylation process in which L-malic acid is converted into L-lactic acid. This process is desirable in most red wines and in some white and sparkling wines for three main reasons: it lowers acidity (leading to a softer mouthfeel), confers microbiological stability, and it modifies the wine’s flavor (Bartowsky, 2017). Several LAB species can perform MLF, with *Oenococcus oeni* being the main responsible in many countries. As such, it is the most extensively studied and widely used as starter culture (Renouf et al., 2006; Nisiotou et al., 2015; Bartowsky, 2017). Recently, *Lactiplantibacillus plantarum* (formerly *Lactobacillus plantarum*) has emerged as an alternative species. It possesses a more diverse enzymatic profile than *O. oeni,* which may significantly influence flavor development (Iorizzo et al. 2016; Krieger-Weber et al., 2020). Additionally, *Lpb. plantarum* has the genetic potential to produce antimicrobial peptides (plantaricins), which could enhance its dominance during MLF and offer the added benefits of reducing the need of sulphurdioxide (SO_2_) commonly used to inhibit undesirable microbial growth (du Toit et al., 2011).

Wine presents a harsh environment due to its high ethanol content, low pH, the presence of SO_2_ and other inhibitory compounds. Not all LAB species or strains can survive under such conditions, but *Lpb. plantarum* has demonstrated stress tolerance comparable to that of *O. oeni* (Iorizzo et al., 2016; Krieger-Weber et al., 2020). Thanks to transcriptomic and other omics-bases analyses, we are beginning to understand the gene and protein expression profiles that contribute to LAB survival in wine (Olguin et al., 2010; 2015; 2019; Bordas et al., 2015; Margalef-Català et al., 2016). However, most of this research has focused on *O. oeni*, while studies involving *Lpb.plantarum* remain limited (Fiocco et al., 2007; Miller et al., 2011).

One of the main criteria for selecting LAB strains as starter cultures is their resistance to wine conditions and their metabolic performance. However, although commercial starter cultures are widely available, a limitation is that they are often derived from specific regions and may not adequately capture the microbial diversity representative of other terroirs. The potential contribution of microorganisms to the concept of terroir has long been overlooked (Knight et al., 2015). These microbes not only participate in fermentation but also influence soil chemistry and nutrition, as well as grapevine health, yield, and quality (Belda et al., 2017). Several studies have linked wine characteristics to regional microbial communities (Vitulo et al., 2019), with some specifically highlighting the role of bacteria in shaping wine quality (Portillo et al., 2016; Grangateau et al., 2017; Lorentzen & Lucas, 2019). Therefore, it is important to evaluate whether isolated microbial strains are beneficial within a given terroir. In some cases, the preservation of terroir itselfmay guide the isolation, identification, and characterization of strains for use as starter cultures.

LAB populations in Patagonian wines display high species and strain diversity during spontaneous MLF (Valdés La Hens et al., 2015). In previous work, we applied an acclimation protocol (Bravo-Ferrada et al., 2014), in which cells were cultured in a broth with moderate ethanol concentration prior to wine inoculation. We investigated the influence of two temperatures (18∟°C and 21∟°C) on gene expression using *O. oeni* strains isolated from Patagonian Pinot noir wines (Olguin et al., 2019). We found that L-malic acid consumption, population survival, andtranscriptional behavior differed depending on acclimation temperature.

The aim of the present study was to evaluate the expression of stress-related genes in *Lpb. plantarum* UNQLp11 induced by acclimation at two temperatures (18∟°C and 21∟°C). Gene selection was based on previous studies involving both *O. oeni* (Olguin et al., 2010; 2019) and *Lpb. plantarum* (Fiocco et al., 2007; Miller et al., 2011). Gene expression was quantified by reverse transcription real-time quantitative PCR (Real Time RT-qPCR). Population survival and L-malic acid consumption after acclimation were monitored in Pinot noir wine.

## Material and methods

### Growth conditions

*Lpb. plantarum* subsp. *plantarum* ATCC 14917 (DSM 20174, Zheng et al., 2020) and the strain UNQLp11 (RGPRN-/-M-/-357-/-Semorile-/-626-/-UNQLp11-/-2024), previously isolated from Patagonian red wines (Bravo-Ferrada et al., 2013; Iglesias et al., 2020) where cultured at 28 °C in tubes containing MRS broth medium (De Man et al., 1960), supplemented with L-malic acid (4 gL^-1^) and fructose (5 gL^-1^) at pH 5.0. After 48 h cells were harvested by centrifugation and resuspended in the acclimation medium (50 gL^-1^ MRS, 40 gL^-1^ fructose, 20 gL^-1^ glucose, 4.5 gL^-1^ L-malate, 1 gL^-1^ Tween 80, 0.1 mgL^-1^ pyridoxine, pH 4.6) containing 6% (v/v) of ethanol (Bravo-Ferrada et al., 2014). One of the acclimation tubes was cultured at 18 °C and the other one at 21 °C. We chose these temperatures because both are common during MLF in wineries. Samples were taken at time zero (at the beginning of cell acclimation) and 48 h after acclimation (just before wine inoculation).

Then, cells were harvested by centrifugation and inoculated in tubes containing sterile Pinot noir wine (vintage 2014), at the final stage of alcoholic fermentation (14.3 % ethanol, pH 3.7, 2.3 gL^-1^ L-malic acid). These tubes were kept at 21 °C and cell survival was analysed every 2-3 days until the 19th day. All assays were performed in duplicate at different times using independent precultures and cell survival was monitored by counting colonies on plates of MRS medium supplemented as described above. L-malic acid consumption was measured using the L-Malic Acid Enology Enzymatic kit (BioSystems SA, Barcelona, Spain) according to the manufacturer’s instructions.

Sovereign Rights Over Natural Resources: The indigenous LAB strain mentioned above is a genetic resources belonging to the Provinces of Rio Negro (UNQLp11). It was deposited in the strain collection of the Laboratorio de Microbiología Molecular (IMBA–DCyT–UNQ-CIC) and registered with the Environment and Climate Change Secretariat— Province of Rio Negro Government. Their preservation follows the protocols of the Convention on Biological Diversity and the Nagoya Protocol, which safeguard the ownership and use of genetic resources from the regions where they were isolated (RESOL-2024-626-E-GDERNE-SAYCC#SGG, NO-2024-00628737-GDERNE-SAYCC#SGG).

### RNA extraction

*Lpb. plantarum* cells were harvested by centrifugation and frozen until RNA extraction. Total RNA extractions were performed using the SV Total RNA Isolation System #Z3100 (Promega) following the manufacturer’s instructions. RNA concentrations were calculated by measuring absorbance at 260 nm using a Nanodrop spectrophotometer (Thermo Fisher Scientific, 1000).

### Selection of candidate genes and nucleotide sequences

The gene sequences of *Lpb. plantarum* reference strain SRCM100442 (Genome assembly ASM991365v1) were obtained from the National Center for Biotechnology Information (NCBI). The reference genome assembly strain SRCM100442 was used for sequence comparison against *O. oeni* PSU-1 (NC_008528) (Table 1) and for primers design when applicable (Table 2). We also tested sequence alignment and *in silico* PCR with the strain ATCC 14917 gene sequences. Sequence alignment and primer pairs were tested in silico using the NCBI Needleman-Wunsch Global Align Nucleotide Sequences and Primer-BLAST, respectively (https://blast.ncbi.nlm.nih.gov/Blast.cgi).

**Table 1.** Comparison of genes sequence alignment between *Lactiplantibacillus plantarum* SRCM100442 and *Oenococcus oeni* PSU-1 (NC_008528)

| Target gene *Lpb. plantarum* | Gene name | Gene sequence identity (%) | Protein sequence identity (%) | *O. oeni* gene symbol |
| --- | --- | --- | --- | --- |
| C7M33_RS14920 | *ldhD* | 60 | 51 | OEOE_RS01985 |
| C7M33_RS06325 | *gyrA* | 66 | 69 | OEOE_RS00030 |
| C7M33_RS14590 | *rpoD* | 59 | 61 | OEOE_RS04785 |
| C7M33_RS01170 | *recA* | 65 | 68 | OEOE_RS06650 |
| C7M33_RS09485 | *gapB* | 72 | 72 | OEOE_RS01940 |
| C7M33_RS05025 | *hsp20* | 57 | 41 | OEOE_RS01385 |
| C7M33_RS11210 | *rfbB (rmlB)* | 57 | 45 | OEOE_RS06990 |
| C7M33_RS14780 | *dnaK* | 70 | 74 | OEOE_RS06310 |
| C7M33_RS09640 | *glmS* | 63 | 60 | OEOE_RS03035 |
| C7M33_RS01055 | *trxA* | 60 | 50 | OEOE_RS08215 |
| C7M33_RS10385 | *cspA* | 79 | 81 | OEOE_RS06620 |
| C7M33_RS04410 | *dacA* | 51 | 32 | OEOE_RS03435 |

Note: *O. oeni* PSU-1 genes locus tag have been discontinued but are still accessible. Since the new sequences are variable, we decided to use the ones we already had to compare with the *Lpb. plantarum* reference genome assembly (strain SRCM100442).

**Table 2.**
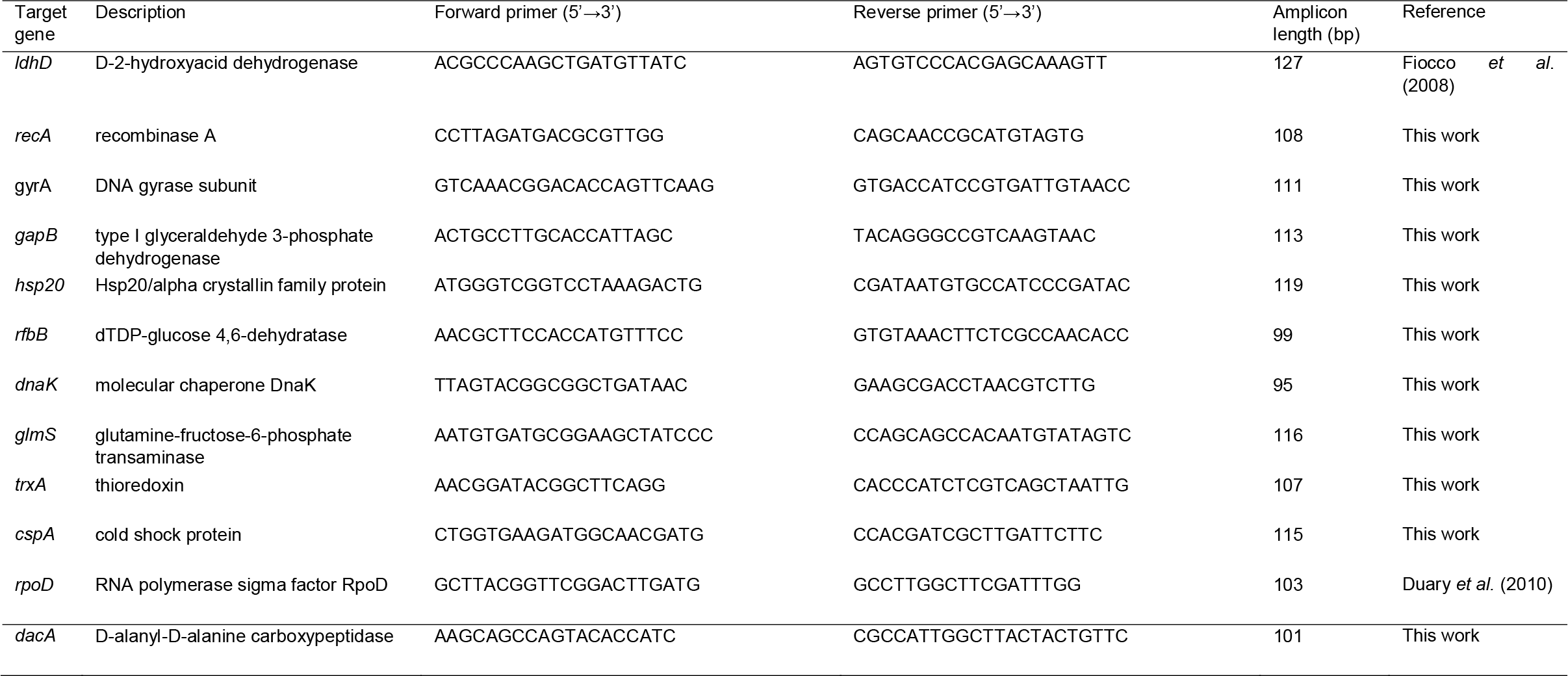
Primers used in this work.

### Reverse transcription and Real Time RT-qPCR

cDNA was synthesized from RNA (15 ng μL^-1^) using TaqMan Reverse Transcription Reagents (Applied Bio-systems) as recommended. Ten pairs of primers were designed (Table 2) to be about 18–22 bases long, to contain over 50 % G/C and to have a melting temperature (Tm) above 60 °C. Two pairs of primers were taken from previous works (Fiocco et al., 2008; Duary et al., 2012). The length of the PCR products ranged from 95 to 127 bp (Table 2). RT-qPCR was performed in 12 μL final volume containing 1.5 ng of total cDNA, 1.0 μL of each primer at 3.15 μmol L^-1^, 2.5 μL of RNase free water and 6.5 μL of SYBR Green PCR Master Mix (Applied Biosystems). Amplifications were carried out using a QuantStudio 3 Real-Time PCR Systems (Applied Biosystems) with an initial step at 95 °C for 5 min followed by 35 cycles of 95 °C for 10 s and 60 °C for 30 s and 72 ºC for 30 s. An additional step to establish a melting curve was used to verify the specificity of the real-time PCR reaction for each primer pair. The threshold value used in this study was automatically determined by the instrument.

Results were analysed using the comparative C_T_ method (2^-ΔΔCT^) as described by Schmittgen & Livak (2008). Two statistical algorithms were used to determine the stability of the candidate reference genes: NormFinder (Andersen et al., 2004) and geNorm (Vandesompele et al., 2002). The three candidates, *ldhD, rpoD* and *gyrA* were chosen as the best combination.

## Results

The *Lpb. plantarum* strain used in this study, isolated from a Patagonian Pinot noir wine undergoing spontaneous MLF, was identified as UNQLp11 (Bravo-Ferrada et al., 2013). It was selected from a pool of strains obtained from the same wine based on its oenological properties (Brizuela *et al*., 2018). *Lpb. plantarum* DSM 20174, isolated from pickled cabbage, was used as a reference strain. Since this strain originates from a non-wine environment, any observed metabolic or genetic similarities are likely species-specific rather than environmentally induced.

Acclimation temperatures of 18∟°C and 21∟°C were selected based on prior work showing temperature-dependent behavior in *O. oeni* strains (Olguin et al., 2019).

During the 48-hour acclimation period, cell density increased by approximately two logarithmicunits (LOGs) at both temperatures for both strains.

Genes from *Lpb. plantarum* were selected based on their known functions, under the hypothesis that they may respond to wine-associated stress conditions similarly to *O. oeni* (Table 1). Prior to quantitative analysis, Real Time RT-qPCR amplification of the selected genes was tested in both strains. The amplified fragment sizes were as expected (Table 2). Amplification efficiency was calculated for each primer pair; the gene *dacA* was excluded due to poor efficiency (E = 80.21%, R^2^ = 0.9958). All remaining primers had efficiencies within 10% of each other, making them suitable for relative quantification using the comparative Ct (ΔΔCt) method (Schmittgen & Livak, 2008).

Relative gene expression was first analyzed using the sample at time zero under acclimation conditions (A0) as the calibrator (Fig. 1). At 18∟°C, the genes *rfbB* (also referred to as *rmlB*) and *glmS* were clearly overexpressed, especially in the Patagonian strain UNQLp11 (Fig. 1A). In contrast, *recA* was slightly overexpressed in the reference strain ATCC 14917 but underexpressed in UNQLp11. Genes *dnaK, trxA, cspA*, and *gapB* were consistently underexpressed in both strainsat this temperature.

**Figure 1.**
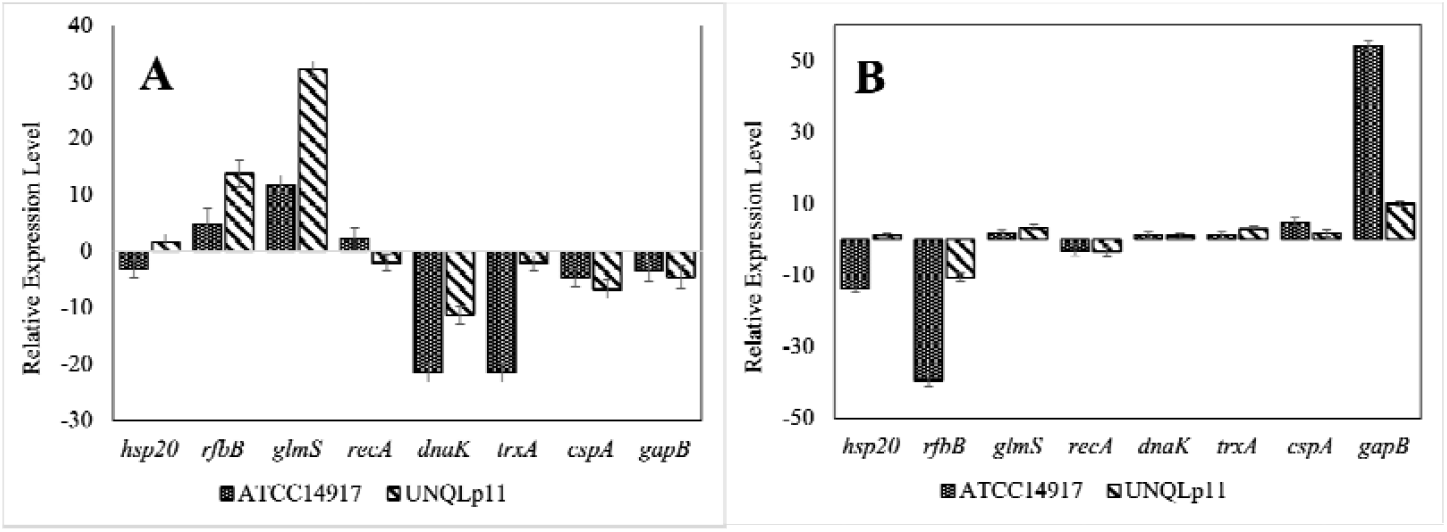
Relative expression of eight genes related to stress response (*hsp20, rfbB, glmS, recA, dnaK, trxA, cspA, gapB*) after 48 h acclimation at 18 °C (A) and at 21 °C (B). The calibration condition used was time zero when *Lactiplantibacillus plantarum* cells were inoculated into the acclimation medium.

At 21∟°C, gene expression trends differed (Fig. 1B). In ATCC 14917, most genes showed stable or reduced expression, except for a slight overexpression of *cspA* and a marked overexpression of *gapB*. In UNQLp11, there was slight overexpression of *glmS* and *trxA*, with *gapB* again notably overexpressed. All other genes were stably expressed, with the exception of *rfbB*, which was significantly underexpressed.

To assess the impact of temperature, gene expression levels after 48 hours of acclimation (A48) were compared between 18∟°C and 21∟°C, using the latter as the calibrator condition (Fig. 2). Expression patterns were largely consistent between strains. Both strains showed strong overexpression of *rfbB* and slight overexpression of *glmS*. Genes *dnaK, trxA, cspA*, and *gapB* were underexpressed under the lower temperature. Differences between strains were limited to slight overexpression of *hsp20* and *recA* in the ATCC 14917 strain.

**Figure 2.**
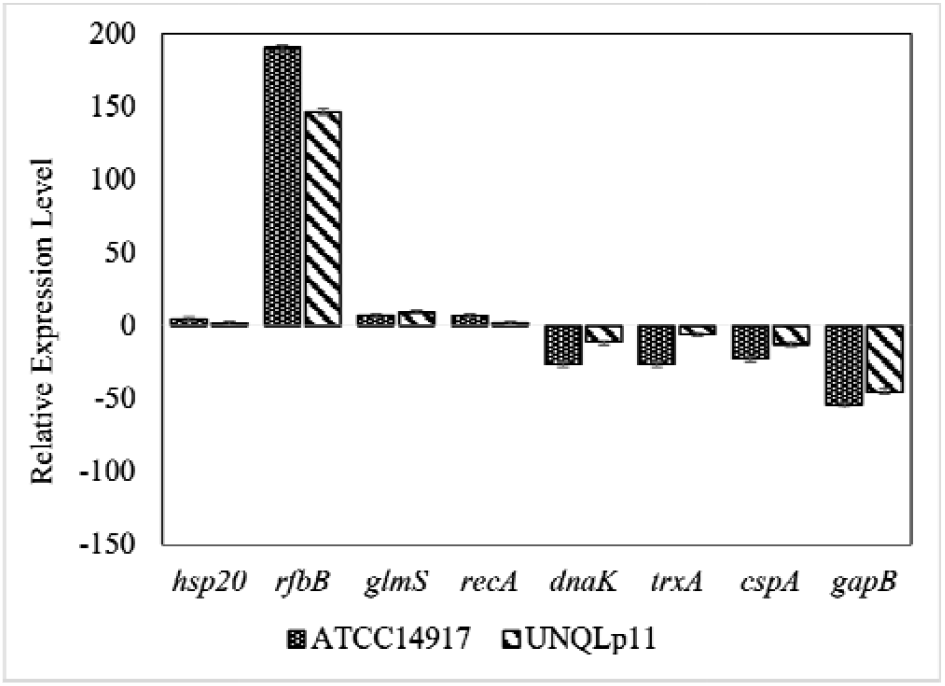
Relative expression of eight genes related to stress response (*hsp20, rfbB, glmS, recA, dnaK, trxA, cspA, gapB*) in *Lactiplantibacillus plantarum* strains after 48 h acclimation at 18 °C. The calibration condition used was time 48 h at 21 ºC.

Cells acclimated at both temperatures (18∟°C and 21∟°C) were inoculated into sterile Pinot noir wine and monitored over 19 days. Cell survival and L-malic acid consumption were evaluated at different time points (Fig. 3).

**Figure 3.**
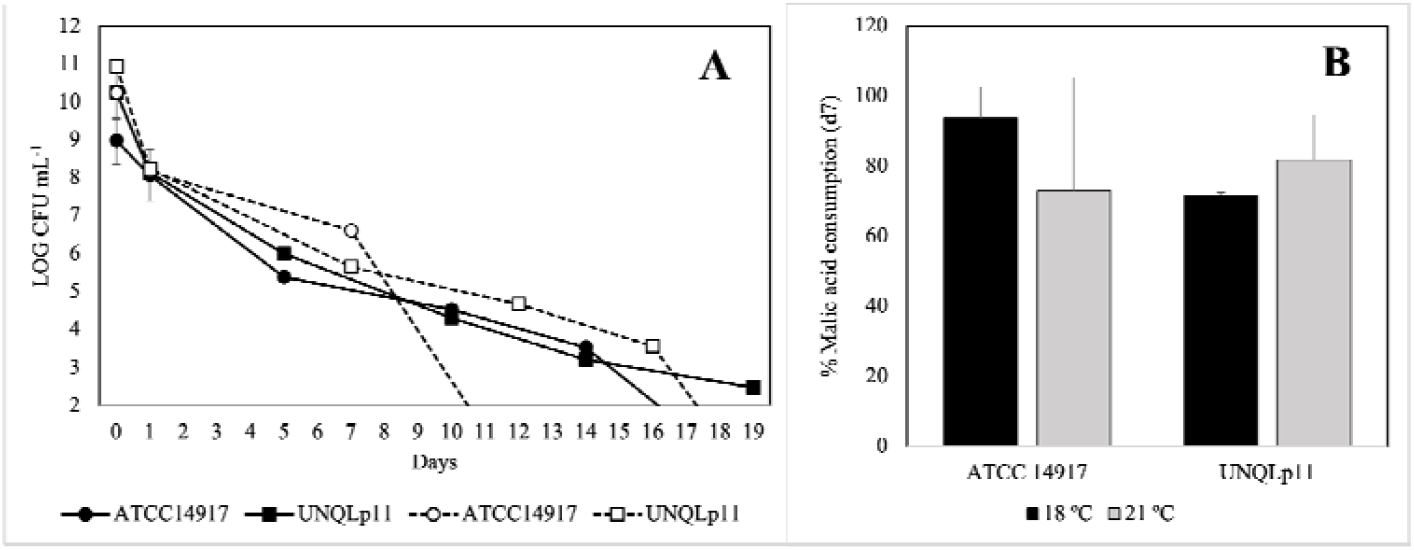
Evolution of cells viability (log CFU mL^-1^) and L-malic acid consumption in Pinot noir wine of the *Lactiplantibacillus plantarum* strains acclimated at 18 and 21 ºC. (A) ATCC 14917 (circles), UNQLp11 (squares), solid symbols and lines indicate strains acclimated at 18 ºC and empty symbols and dashed lines indicate strains acclimated at 21 ºC. (B) Percentage of L-malic acid consumed by cells acclimated at 18 ºC (black bars) and at 21 ºC (grey bars) after 7 days (7d) in sterile wine.

Viable cell counts for both ATCC 14917 and UNQLp11 declined during the fermentation period (Fig. 3A). However, both strains were capable of consuming the majority of the L-malic acid within the first seven days of incubation (Fig. 3B).

## Discussion

The aim of this study was to expand our understanding of the genetic mechanisms involved in the acclimation of *Lpb. plantarum*, which may enhance its survival under the stressful conditions present in wine. Previous studies have demonstrated that wine-associated LAB, such as *O. oeni* and *Lpb. plantarum*, benefit from an acclimation step prior to inoculation into wine (Cecconi et al., 2009; Bravo-Ferrada et al., 2013; Costantini et al., 2015). Stress-related genes in *O. oeni* have been proposed as markers of cellular adaptation (Hamon et al., 2013), but the genetic response of *Lpb. plantarum* to wine-like conditions remains poorly characterized. Thus, we evaluated the expression of stress-related genes in two *Lpb. plantarum* strains subjected toacclimation at two different temperatures.

Sequence comparisons between *O. oeni* and *Lpb. plantarum* indicate that while their genes and proteins perform homologous functions, their regulatory mechanisms can differ. It is known that different stress-resistance phenotypes may protect against the same environmental stressor, even when the molecular pathways involved are distinct (Papadimitriou et al., 2016). Therefore, it is not unexpected that some genes in *Lpb. plantarum* respond similarly to those in *O. oeni*, while others do not (Fig. 1).

The gene expression profiles of strains UNQLp11 and ATCC 14917 showed broadly similar responses, which corresponded with their survival and L-malic acid consumption in wine following acclimation. Among the stress response genes (*hsp20, cspA, dnaK*, and *trxA*), we observed general under-expression at 18∟°C. These temperature conditions are moderate, and more extreme values are typically required to elicit strong responses in heat-shock or cold-shock genes (Liao et al., 2010; Fiocco et al., 2007). Nevertheless, due to the potential for cross-protection mechanisms (Papadimitriou et al., 2016; Ma et al., 2021), we included both *hsp20* and *cspA* in our analysis. The under-expression of *cspA* may align with previous findings in *O. oeni*, where theprotein was downregulated following acclimation in 8% ethanol (Cecconi et al., 2009a).

While earlier studies reported that *Lpb. plantarum* strains overexpressing heat shock proteins were more ethanol-tolerant (Fiocco et al., 2007), our results differ from some previous findings. For example, *hsp* gene expression in *Lpb. plantarum* has been correlated with that in *O. oeni* (Capozzi et al., 2012), but in our study, *hsp20* did not show significant upregulation, consistent with previous work on *O. oeni* under similar conditions (Olguin et al., 2019). Similarly, although *dnaK* expression has been shown to increase under various stress conditions (Liao et al., 2010; Reverón et al., 2012), we did not observe this in our study, again aligning with earlier findings in *O. oeni* (Olguin et al., 2019). The gene *trxA*, typically upregulated under oxidative and ethanol stress in both species (Serrano et al., 2007; Reverón et al., 2012; Margalef-Català et al.,2017), also did not show the expected increase in our experiment.

With respect to carbohydrate metabolism, *gapB* is often considered a housekeeping gene (Duray et al., 2012), yet in our study, its expression appeared temperature-dependent. This aligns with previous observations where *gapB* was upregulated under oxidative stress in specific *Lpb. plantarum* strains (Serrano et al., 2007). Additionally, other studies have reported changes in protein expression linked to carbohydrate metabolism in response to different culture conditions in both *Lpb. plantarum* (Cecconi et al., 2009b; Hamon et al., 2013) and *O. oeni* (Cecconi et al.,2009a; Olguin et al., 2015; Costantini et al., 2015).

In the context of amino acid metabolism, we analyzed *glmS*, which showed a notable increase in expression, especially at 18∟°C. Previous work has shown that *glmS* is downregulated in response to antioxidant stress induced by hydroxytyrosol or gallic acid (Reverón et al., 2015; 2020), and its expression also decreases in *O. oeni* following ethanol shock (Olguin et al., 2015). Our results contrast with those findings, suggesting a different regulatory pattern undertemperature-induced stress.

The *recA* gene, associated with DNA repair and recombination, has been variously classified as a housekeeping or stress-related gene (Miller & Kokjohn, 1990; Curiel et al., 2011; Reverón etal., 2012, 2015). In our study, its expression remained relatively stable under both conditions.

An especially interesting result was observed for *rfbB*, a gene involved in cell envelope biogenesis. It was strongly overexpressed at 18∟°C but downregulated at 21∟°C. Previous studies have reported no significant change in *rfbB* protein levels under acidic conditions in *Lpb. plantarum* (Hamon et al., 2013), while in *O. oeni*, both over- and under-expression have beenobserved depending on the acclimation conditions (Olguin et al., 2015; 2019).

The impact of acclimation on both *O. oeni* and *Lpb. plantarum* using the same medium has been studied in previous research (Bravo-Ferrada et al., 2014; Brizuela et al., 2018). Thus, the similar survival and L-malic acid consumption profiles observed in both *Lpb. plantarum* strains in this study were not unexpected. Notably, strain UNQLp11 was isolated from Patagonian Pinot noir wine, and the selection of indigenous strains as starter cultures is gaining global interest for enhancing regional wine identity (Bartowsky et al., 2017). As suggested by Mao et al. (2021), environmental selection pressure can drive genomic changes that improve strain adaptability to specific ecological niches. Therefore, while our results are promising, further research usingadditional wine-isolated *Lpb. plantarum*strains is necessary to validate these findings.

To our knowledge, this is the first study to evaluate stress-related gene expression in *Lpb. plantarum* as potential markers for winemaking applications. As the use of this species in starter culture development continues to grow, further research incorporating gene expression and omics approaches will be valuable. Similar to ongoing work in *O. oeni*, these studies can help correlate genetic behavior with enological performance, ultimately supporting the selection of robust, regionally adapted strains for the wine industry.

In conclusion, this study offers new insights into the genetic mechanisms underlying the acclimation of *Lpb. plantarum* to wine conditions—an area that remains less understood compared to *O. oeni*. By analyzing the gene expression profiles of *Lpb. plantarum* strains at two temperatures, we identified both shared and distinct responses relative to *O. oeni*. While several stress-related genes (*hsp20, cspA, dnaK, trxA*) were under-expressed, genes suchas *glmS* and *rfbB* showed temperature-dependent overexpression.

These findings highlight the potential of *Lpb. plantarum* as a viable candidate for use in starter culture development, particularly when selecting indigenous strains to support regional wine identity. Overall, the results deepen our understanding of *Lpb. plantarum*’s stress adaptation and support its further exploration for enological applications, laying the groundwork for future studies aimed at optimizing strain selection and performance in winemaking.

## Acknowledgements

We would like to thank the Environment and Climate Change Secretariat, Government of the Province of Rio Negro, for authorizing access to provincial genetic resources and their use for academic and research purposes. The authors also thank the Laboratorio de Microbiología Molecular (LMM) of the Universidad Nacional de Quilmes (UNQ) for lending them the equipment to perform the Real Time RT-qPCR.

## Data information availability

Data are available online: https://doi.org/10.1101/2025.08.28.672453; Zerbino et al., 2025. Additional documents are available: http://hdl.handle.net/11336/270401; Zerbino et al., 2025.

